# Running the distance: embodied distance measurement is robust to social interference in fiddler crabs

**DOI:** 10.1101/2025.07.27.666470

**Authors:** Hisashi Murakami, Takenori Tomaru

## Abstract

Path integration (PI) is an essential navigation strategy. Through PI, animals integrate their movements during foraging into a ‘‘home vector” that allows direct return to the origin. In humans, feedback from body movement contributes to readout of the home vector: physical estimation by walking toward the origin improves people’s ability to identify the origin point. However, how such feedback affects home vector readout in other animals remains unclear. Here, we show that leg movement feedback contributes to a robust readout of the home vector despite social interference in fiddler crabs. We found that whereas the internal estimation of burrow location during the return path is subject to induced errors by social behavior, the physical estimation is not, and feedback from leg movement contributes to robust distance measurement. Our findings suggest that animals may have evolved complex navigational systems to handle the inevitable interplay between navigational cues and social behavior.

## Introduction

Many animals, including humans, use a spatial memory system for navigation known as path integration (PI), through which an animal’s movements are summed to form a single “home vector” that enables direct return to the origin point [1, 2]. When measuring distance, animals generally use some sort of idiothetic sensory information [3–6], i.e., information created by and derived from an animal’s own locomotion. In humans, walking itself can be the source of information not only when measuring distance during outbound journeys (i.e., forming the home vector) but also when returning to the origin point (i.e., reading it out) [7–9], suggesting that PI is an embodied memory process [10]. Moreover, physical estimation by walking toward the origin improves people’s ability to identify the origin point [9]. Although the home vector can be read out without feedback from body movement, such feedback contributes to its readout for an effective return.

In other animals, the role of feedback from body movement in reading out the home vector remains unclear. In desert ants and fiddler crabs, stride integration is known to be used for distance measurement [3, 4, 11]. By integrating the length of the steps taken, these animals can determine the distance traveled during the return path. They are suggested to compare this with the desired travel distance (the length of the home vector) to assess whether they have reached the origin point [12, 13]. In this view, leg movements are considered crucial to get the distance traveled, whereas the desired travel distance is assumed to be always accessible as a kind of internal representation, independent of body movements.

Yet, body movement feedback might also play a role in accessing the desired travel distance (that is, reading out the length of the home vector). For example, these animals sometimes must access PI memory to internally estimate the origin point (e.g., location of nest or burrow) partway through the return path, when comparing and interacting with other information, such as visual cues [14–17]. Could physical estimation by walking toward the origin improve their ability to identify the origin point compared to this internal estimation partway through the return path? If so, body movement feedback could contribute to reading out the home vector. Specifically, this feedback contributes not only to monitoring the traveled distance during the return (via stride integration) but also to reading out the desired travel distance. However, it is hard to compare the accuracy of these two processes because of the difficulty of directly assessing the accuracy of internal estimation in nonhuman animals. One alternative would be to investigate situations that affect these two processes differently, e.g., where an error occurs in the internal estimation partway through the return path but not in the physical estimation.

Fiddler crabs are potentially suitable for studying this difference, as their frequent engagement in social behaviors likely introduces errors into PI [18, 19]. When threatened, these crabs can run back to their own burrows precisely despite generally lacking visual contact with the burrows because of their limited perspective [20–24]. They traditionally have been considered to do this by relying exclusively on PI (e.g., [20, 24]), irrespective of the presence of landmarks near their burrows, suggesting that the visual mechanism basically does not function during a rapid escape run. However, in our previous experiments, we observed that the fiddler crab *Austruca perplexa* can respond to visual cues even during escape runs when PI and visual cues are in conflict [25, 26]. We masked true burrows (the goal of PI), and placed fake entrances (visual cues) only along their homing paths, given that *A. perplexa* can fully perceive a burrow entrance, regardless of body size, only when it is within about 4 cm [27, 28]. We found that threatened fiddler crabs only responded to visual cues located near their burrows; otherwise, they ran up to the masked true burrow, ignoring the fake entrance [26]. This demonstrates that crabs exhibit a clear transition in behavior during escape runs: they initially suppress visual cues in favor of PI memory, but then switch to prioritizing visual cues as they approach their burrow.

In this study, to explore whether and how social behavior influences PI and its interaction with visual cues, we observed escape responses of *A. perplexa* that started during territorial behavior (Fig. 1). Adopting our previous approach [26], we additionally placed a conspecific contained in a transparent cup 18 cm from the focal crab’s burrow and located a fake entrance along the homing path of the focal crab engaged in territorial behavior (Methods). Under this social condition, we found that the clear transition observed in the non-social condition in our previous study [26] was disrupted. This suggests that error was induced by social behavior into the internal estimation of burrow locations, which is used in interactions with visual cues during homing runs. Nevertheless, we also confirmed that, when crabs did not respond to the fake entrance, they could return to the masked burrow location just as precisely under the social condition as under the non-social condition. These results suggest that the internal estimation of burrow location during the return path is subject to induced errors, but the physical estimation is not, and that feedback from leg movement contributes to robust distance measurement.

**Figure 1.**
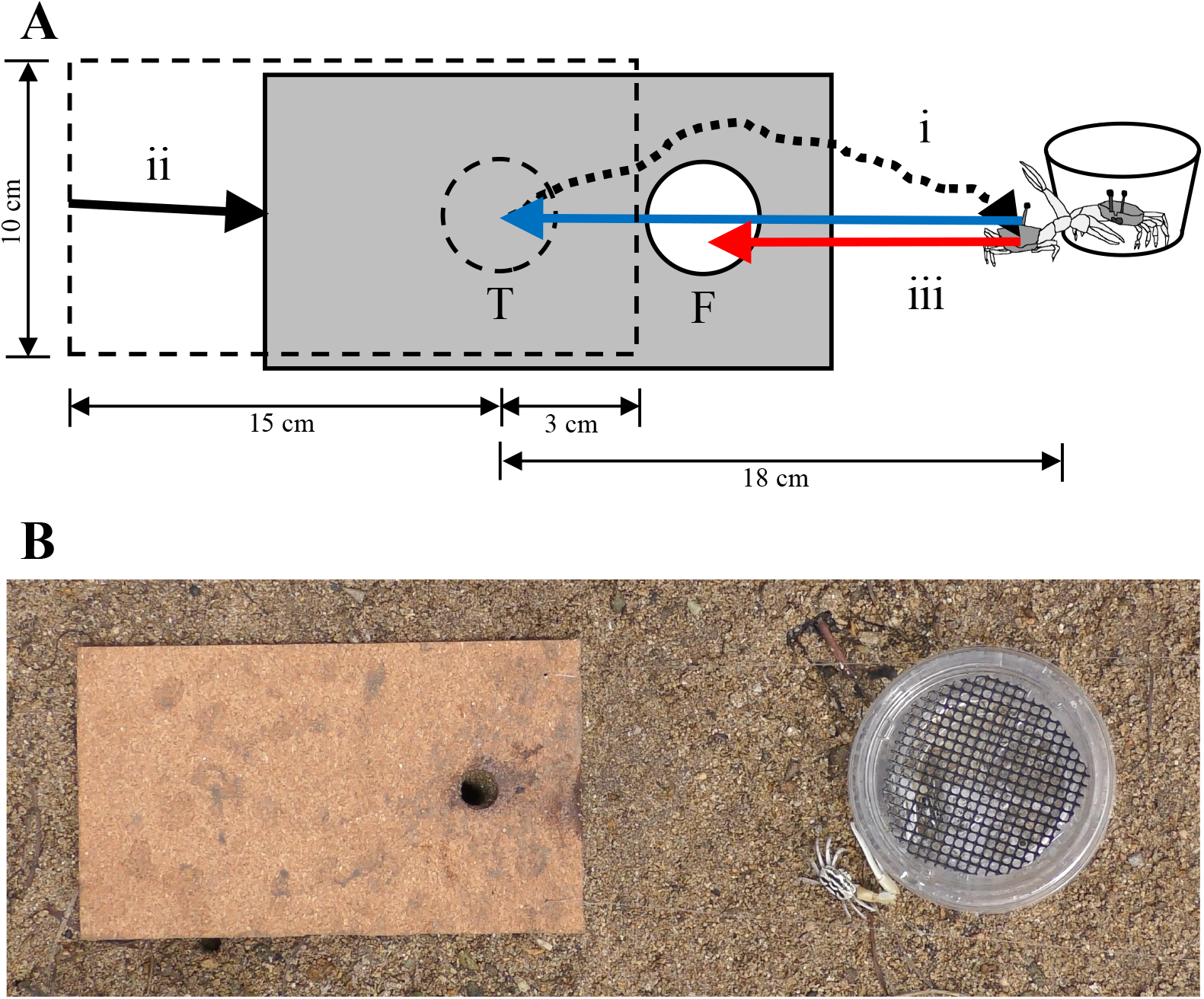
Experimental paradigm. **(A)** Illustration of the experimental procedure. After the focal crab had emerged from its own true burrow (on which a fake entrance cut into a cork sheet was superimposed) and moved toward a transparent cup containing a conspecific to engage in territorial behavior (i), the sheet was translocated so that the true burrow was masked and only the fake entrance was shown (ii). The focal crab ran back to the masked true burrow or fake entrance when it was startled (iii; blue or red arrow, respectively). Dashed circle (T): true burrow; solid circle (F): fake entrance. **(B)** A snapshot from the experiment showing the crab just after leaving its burrow and approaching a conspecific, immediately before the sheet was moved.

## Results

After being threatened while engaging in territorial behavior, fiddler crabs rushed back and stopped abruptly at the fake entrance (“F event,” Methods and Movie S1) or the masked true burrow (“T event,” Movie S2). In Fig. 2A, we show the distribution of fake entrance locations during F and T events under the social condition, in a reference frame where we defined the positions of the true burrows as the origin and situated the direction from the starting point of the run to the true burrow to be from right to left along the *x*-axis. We observed that the distributions of the T and F events under the social condition (the present study; Fig. 2A) overlap more compared to the non-social condition (our previous study [26]; Fig. S1A). We then calculated the probability of occurrence of a T event in relation to the distance between the true burrow and the fake entrance along the *x*-axis, as well as the curve fitted by sigmoid functions. Figure 2B shows the data and fitted curve under the social condition, with the fitted curve under the non-social condition overlaid for comparison (see also Fig. S1B for the non-social data points). If social behavior alters the priorities of PI and visual cues and makes crabs return to fake entrances more often even at larger distances, the threshold of the logistic curve (i.e., the distance of 50% probability) should be larger under the social condition. However, there was no significant difference in the threshold between conditions (social: 44.18 mm; non-social: 43.43 mm, Bootstrap test, *p* = 0.854). Rather, the social condition had a significantly shallower slope than under the non-social condition (0.075 vs. 0.20, Bootstrap test, *p* = 0.008). These results suggest that social behaviors disrupt the clear transition observed in the non-social condition, i.e., from initially suppressing visual cues in favor of PI memory to later prioritizing them. Taking previous research into account [18, 19], social behaviors seem to introduce errors in PI (i.e., the internal estimation of PI goal (the true burrow location), which interacts with visual cues).

**Figure 2.**
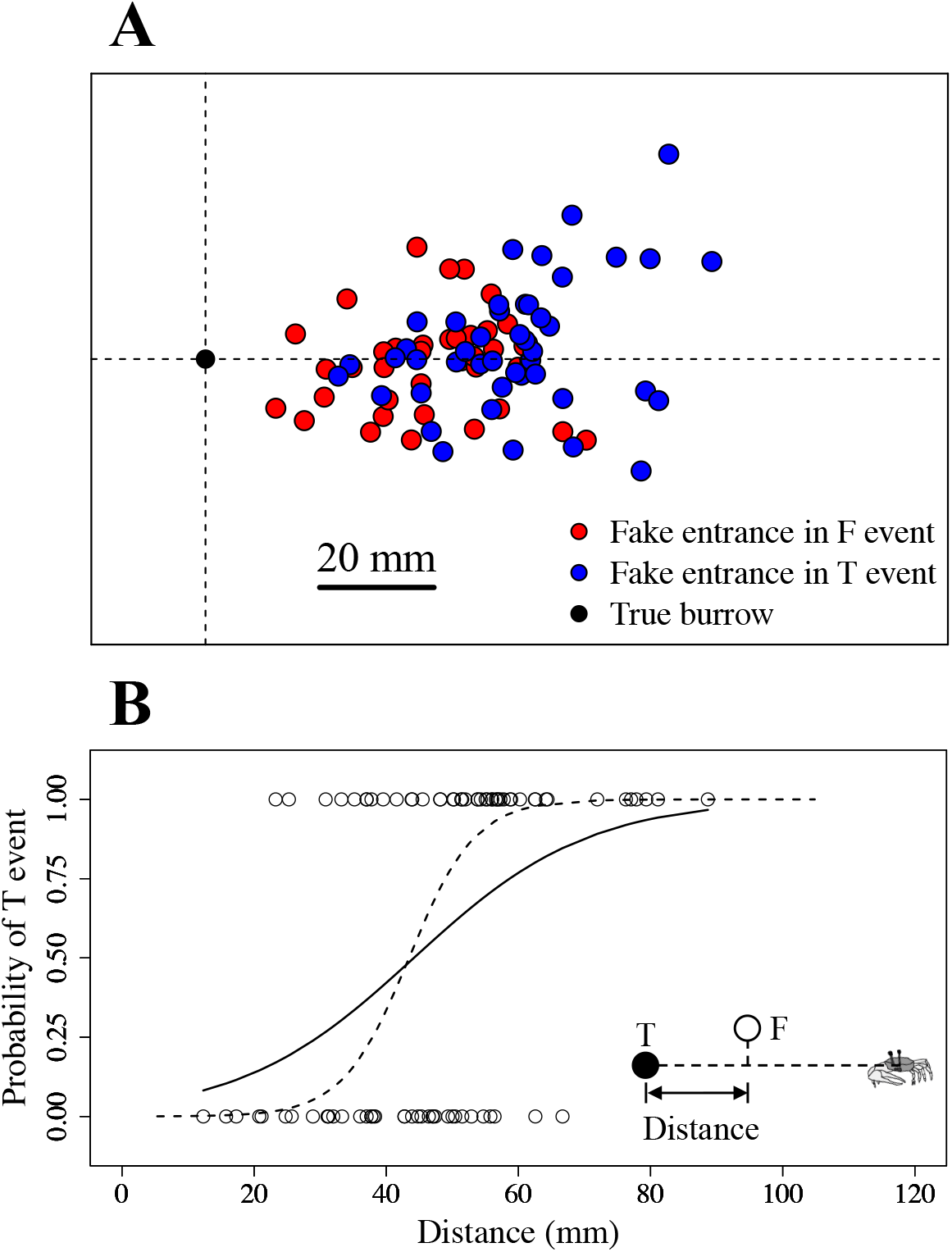
Burrow selection in relation to the distance between the true burrow and fake entrance under the social condition (experiments with territorial behavior). **(A)** Location distribution of fake entrances during F events (returned to fake entrance, red circles) and T events (returned to true burrow, blue circles). The black circle indicates the true burrow location. The direction from the starting point of the run to the true burrow is normalized to be right to left. **(B)** Probability of occurrence of a T event in relation to the distance between the true burrow and fake entrance. Open circles represent data from individual trials: T events (1) and F events (0). The solid line represents the best-fit sigmoid function. The dashed line shows the sigmoid curve under the non-social condition (experiments without territorial behavior), adopted from ref. [26] for comparison. The inset illustrates how the distance between the true burrow and the fake entrance is determined (see main text for details). See also Fig. S1 for data points under the non-social condition.

How much error does social behavior introduce into the internal estimation of the true burrow location? To examine this, we conducted a simulation using data from the non-social condition, where we added artificial errors to the location of true burrows (the goal of PI), with the errors normally distributed with various values of standard deviation (*σ*). When we calculated the slope of the fitted sigmoid curve using the *x*-distance between this modified burrow and the fake entrance for each *σ*, we identified that the slope with a *σ* of 18 mm closely matches that of the social condition (see Fig. 3 and Methods).

**Figure 3.**
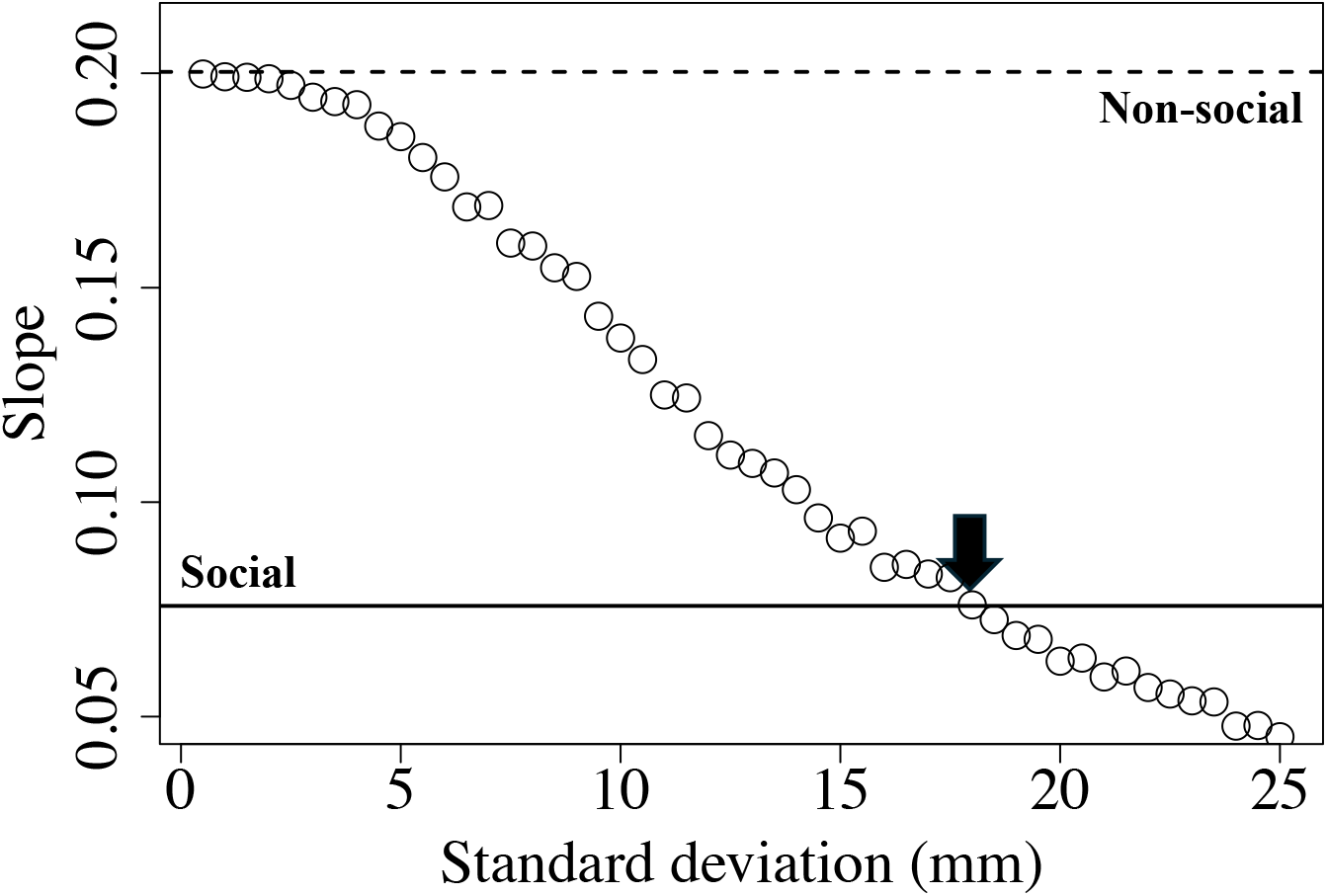
Results of a simulation to estimate how much error social behavior introduced in PI memory. We virtually displaced the true burrow locations under the non-social condition by using normally distributed errors with various values of standard deviation (*σ*). The slope of the fitted sigmoid curve using these modified burrow locations was plotted as a function of *σ*. The slope with *σ* set to 18 mm (black arrow) closely matched the data obtained under the social condition. The dashed and solid horizontal lines represent the slope under the non-social and social condition, respectively.

Does social behavior also induce error when crabs physically run up to the masked true burrows? To address this, we determined “homing error” during T events (returned to masked true burrow) by calculating the distance between the position of the masked true burrow and the crab’s position when it stopped. If social behavior also induces error in the physical estimation, this homing error should be larger under the social condition than under the non-social condition. However, we found no significant difference in homing error between the social and non-social conditions (Welch’s *t*-test, *N*_*social*_ = 46, *N*_*non-social*_ = 25, *t* = 0.29, *p* = 0.77, Cohen’s *d* = 0.068) (Fig. 4). Moreover, we estimated a “predicted error” by adding Gaussian noise with the estimated *σ* of 18 mm to the true burrow position of the non-social data and calculating the distance between this simulated true burrow location and the crab’s position when it stopped (Methods). We then observed that the homing error under the social condition was significantly smaller than this predicted error (Welch’s *t*-test, *N*_*social*_ = 46, *N*_*predicted*_ = 25, *t* = −4.42, *p* = 0.00023, Cohen’s *d* = −1.32) (Fig. 4). This implies that the crabs successfully returned to the masked true burrow even under the social condition, and that social behavior does not influence homing error in the physical estimation while running to the burrow. Taken together, these results suggest that the two readout processes of the home vector are differently influenced by social behavior: the internal estimation of the burrow location during the return path is subject to induced errors, but the physical estimation is not.

**Figure 4.**
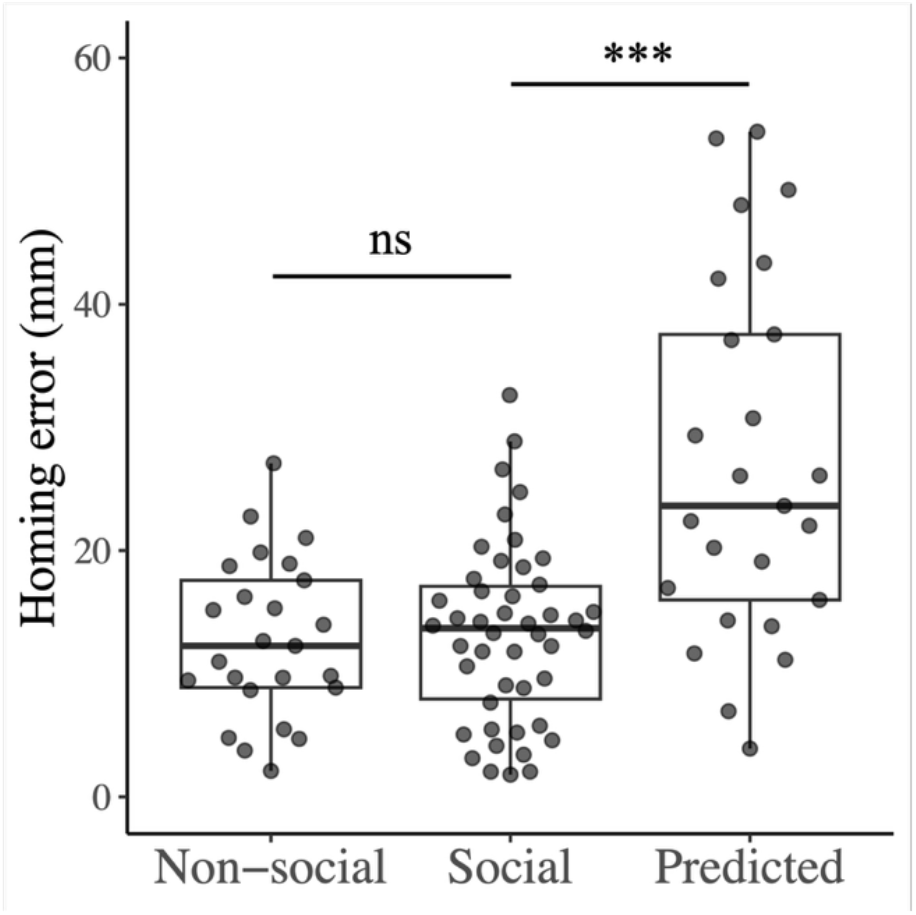
Homing error during T events (return to masked true burrow). Homing error was calculated as the distance between the position of the masked true burrow and the crab’s position when it stopped. Homing error in the social condition (center) did not differ significantly from that in the non-social condition (left, *p* > 0.05) but was significantly smaller than the predicted error (right, *p* < 0.001). Predicted error was calculated by adding Gaussian noise with an estimated standard deviation of 18 mm to the true burrow positions from non-social data and calculating the homing error with this modified burrow location. Each data point represents a trial. Box-and-whisker plots show the median (thick central line), interquartile range (box), and 1.5× the interquartile range (whiskers). ns, not significant.

## Discussion

In this study, we investigated whether and how social behavior influences PI and its interaction with visual cues. To this end, we observed the escape responses of fiddler crabs, *A. perplexa*, in experiments that set PI and visual cues in conflict by masking true burrows (the goal of PI) and placing fake entrances (visual cues) along their homing paths. After being threatened, fiddler crabs ran back and stopped suddenly at the fake entrance (F event) or the masked true burrow (T event). Their responses to visual cues differed depending on whether they engaged in territorial behavior before the homing runs (i.e., social condition or non-social condition [26]).

To quantify this, we calculated the probability of a T event in relation to the true burrow–fake entrance distance (Figs. 2 and S1). Under the non-social condition, the fitted sigmoid curve had a steep slope, indicating a clear transition from the prioritization of PI to visual cues during the homing run (Fig. S1B). If social behavior altered the priorities of PI and visual cues, such that crabs prioritized visual cues more often even at larger distances, then the threshold of the sigmoid curve should have shifted to larger values under the social condition. However, the threshold did not differ between conditions. Instead, the slope of the curve became shallower (Fig. 2B). Building on previous studies suggesting that social behavior introduces errors into PI [18, 19], these results can be interpreted as indicating that social behavior imposed errors on the PI goal (the internal estimation of the true burrow location, which interacts with visual cues). A simulation with the non-social data in which Gaussian errors were added to the true burrow locations (PI goals) showed that a standard deviation of 18 mm reproduced the same slope observed under the social condition (Fig. 3). Interestingly, when crabs physically run up to the masked true burrows (T event), homing error did not differ between the social and non-social condition (Fig. 4). These results suggest that the internal estimation of burrow location during the return is subject to induced errors, whereas the physical estimation is not.

As has been well documented, fiddler crabs can start their homing run even when they cannot see their burrow [20–24]. This requires crabs to internally estimate their burrow location even in the absence of body movement. In other words, the home vector can be read out without body movement as some sort of internal representation that signals the location of the burrow before initiating the homing run. Perhaps this internal estimation is also used to compare and interact with visual cues, i.e., to determine whether to respond to these cues, and social behavior induces error into it. On the other hand, information from leg movements can also contribute to reading out the home vector, and this information is presumably primarily used to assess when to stop running and is not influenced by social behavior. Hence, when crabs ignore visual cues (i.e., T events), they can physically run up just as precisely to the masked true burrow even after engaging in territorial behavior. In other words, if feedback from leg movements plays no role in determining when to stop, and if that determination relies solely on the internal representation, the homing error under the social condition would be expected to be larger than that under the non-social condition. Yet this was not the case.

During homing runs, fiddler crabs appear to measure the traveled distance by integrating the length of strides taken, as has also been observed in desert ants [3, 4], rather than by using linear acceleration or optic flow [11]. Our results suggest that feedback from leg movement also contributes to accessing the strides to be taken or the desired travel distance, which is compared with the traveled distance to determine when to stop. In other words, the desired travel distance (the length of the home vector) becomes apparent only when it is exhausted; otherwise, it is not directly available and must be estimated, for example, when comparing with other information (e.g., visual cues).

Fiddler crabs exhibit not only striking navigational abilities during homing runs when threatened, but also various social behaviors such as territorial and courtship behaviors, where male crabs use their greatly enlarged claw to compete with others and attract females (e.g., [29, 30]). This suggests that spatial cognition and social behavior are inherently intertwined. Errors induced by social behavior may require flexibility in spatial cognition, which in turn may allow animals more freedom to engage in social activities. We believe that our findings provide insights into how animals have evolved complex navigational systems to handle the inevitable interplay between navigational cues and social behavior.

## Methods

The present experiment (i.e., the social condition) was conducted in August 2023 during the daytime from 2 h before to 2 h after low tide on a colony of fiddler crabs (*Austruca perplexa*; formerly *Uca perplexa*) on a tidal flat on Iriomote Island, Okinawa, Japan (24°24′N, 123°48′E). Note that this experiment was carried out at the same location and during the same season as our previous experiment (the non-social condition) [26]. We recorded all experiments from above by using a Panasonic HC-WX990M-K camcorder (3840 × 2160 pixels, 30 frames per second) fixed to a tripod. Immediately after the camera was set up, we conducted each experimental trial on an arbitrarily chosen individual in the colony with up to 4 repetitions.

### Experimental Procedure

The procedure for the present experiment (the social condition) was adopted from our previous experiment (the non-social condition) [26], with some minor revisions to introduce territorial behavior between a focal and conspecific male (Fig. 1). We created a fake entrance by cutting a 1.5-cm-diameter hole at a distance of 3 cm from the edge of a 10 × 18 cm cork sheet, to which a plastic sheet of the same size was attached for reinforcement (total thickness, 2 mm). The sheet was attached to a fishing line on the two corners closest to the fake entrance, allowing an observer to pull the sheet towards them.

Before starting trials with each focal male, we captured another male with a burrow at least 2 m away from the focal male’s burrow and placed it in a transparent plastic cup (base diameter 7 cm, height 4.5 cm) with 2–3 mm of water to display it as a decoy in the trials.

Before starting each trial, the fake entrance was superimposed on the true burrow entrance with the focal crab in it. We observed that the crabs emerged from this modified burrow entrance, foraged, engaged in homing behavior, and defended the burrow in a similar manner to what is observed under natural conditions. We placed the transparent plastic cup containing a conspecific male 18 cm from the true burrow, on the shorter side of the sheet from the fake entrance. After the focal crab emerged from and left its burrow, it immediately approached the decoy crab in most cases and engaged in territorial behavior, which involved touching its claws to the cup containing the decoy crab (Fig. 1(i)). An observer, sitting on a chair 2 m away, then translocated the sheet to allow the fake entrance to be shown at various distances along the homing path of the focal crab and the true entrance to be covered by the sheet (Fig. 1(ii)). The sheet translocation was performed slowly to avoid disturbing the focal crab. Then, the focal crab was startled by the observer getting up from the chair, and it ran back to the masked true burrow or to the fake entrance (Fig. 1(iii)).

In total, 94 trials were conducted with 50 crabs. In 10 of the 94 trials, the crab was caught by the edge of the sheet; these trials were excluded. The remaining 84 trials (involving 47 crabs) were used in the following analysis.

Note that our previous experiment (the non-social condition) was conducted without territorial behavior, i.e., without a cup containing a conspecific (80 trials with 58 crabs; see ref. [26] for further details). In this experiment, after the crab emerged from and left its burrow to feed, the sheet was translocated to allow the fake entrance to be shown at various distances along the homing path of the focal crab and the true entrance to be covered. The other procedures were the same as in the present experiment.

### Data analysis

From the recorded footage, we recorded the start and end points of an escape run as well as the locations of the true burrow and of the fake entrance after translocation by using image-processing software (Library Move-tr/2D ver. 8.31; Library Co. Ltd., Tokyo, Japan). We used a reference frame in which the location of the true burrow was defined as the origin, and the direction from the starting point of the run to the true burrow was normalized to be right-to-left along the *x*-axis. We first checked if the fake entrance was translocated outside the area where the crab could fully see it, that is, the area within 4 cm of the home vector (i.e., whether the absolute value of the *y*-coordinate of the fake entrance was smaller than 4 cm in the reference frame described above). This was done because the visual contact distance of *A. perplexa* to its burrow is limited owing to perspective foreshortening, so that the crab can fully perceive the burrow entrance, regardless of its body size, only when it is within about 4 cm [26, 27, 28]. We confirmed that in all 84 trials, the fake entrance was translocated within this area.

If the stopping point was closer to the fake entrance than to the masked true entrance, we considered that the crab had reverted to the fake entrance and labelled these trials “F events.” Alternatively, when the stopping point was closer to the masked true entrance than to the fake entrance, we considered that the crab had run back to the true burrow and labelled these trials “T events.”

We calculated a sigmoid function obtained from logistic regression to evaluate the relationship between the probability of a T event, *P*_*T*_, and the distance between the true burrow and the fake entrance along the *x*-axis (the *x*-coordinate of the fake entrance in the reference frame described above), *D*,

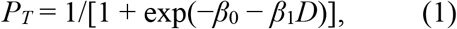

where *β*_0_ and *β*_1_ are the intercept and steepness parameters, respectively.

To estimate how much error social behavior introduces to PI, we conducted a simulation by using data from the non-social condition [26], where we virtually displaced the true burrow locations (i.e., the origin in the reference frame described above) by using normally distributed errors with various values of standard deviation (*σ*). As a result, the simulated coordinate of the true burrow (*x*_*sim*_, *y*_*sim*_) was calculated as

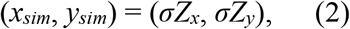

where *Z*_*x*_ and *Z*_*y*_ are Gaussian random numbers with mean 0 and standard deviation 1. Using this simulated coordinate of the true burrow and its distance to the fake entrance along the *x*-axis, we calculated the slope of the fitted curve for each *σ* ranging from 0.5 to 25 mm (in increments of 0.5 mm), averaged across 100 simulation trials, to identify which value of *σ* matched the data observed under the social condition (Fig. 3).

### Statistical analysis

To compare the threshold (−*β*_0_/*β*_1_) and slope (*β*_1_) of the fitted logistic regression models between conditions, bootstrap resampling (1000 iterations) was performed, and *p* values were computed by doubling the proportion of samples in which the estimated difference between conditions crossed zero. Welch’s *t*-tests were used to test differences of homing error between the social condition, non-social condition, and predicted error. When making multiple comparisons, *p* values were adjusted by using the false discovery rate control [31]. All statistical analyses were conducted in R version 4.4.1 (The R Foundation for Statistical Computing, Vienna, Austria).

## Acknowledgements

We are grateful to Iriomote Station, Tropical Biosphere Research Center, University of the Ryukyus, for its kind support of our study, and to Hitonaru Nishie for his helpful comments on earlier versions of the manuscript. This work was supported by JSPS KAKENHI grants [numbers JP18K18348 and JP25K22819].

## Author contributions

H.M. conceived the study, analyzed the data, and wrote the first manuscript draft; H.M. and T.T designed and performed the experiments and contributed to writing the final version.

## Data and materials availability

All data needed to evaluate the conclusions of the paper are present in the paper and/or the Supplementary Materials.

## Supplementary materials

**Figure S1.**
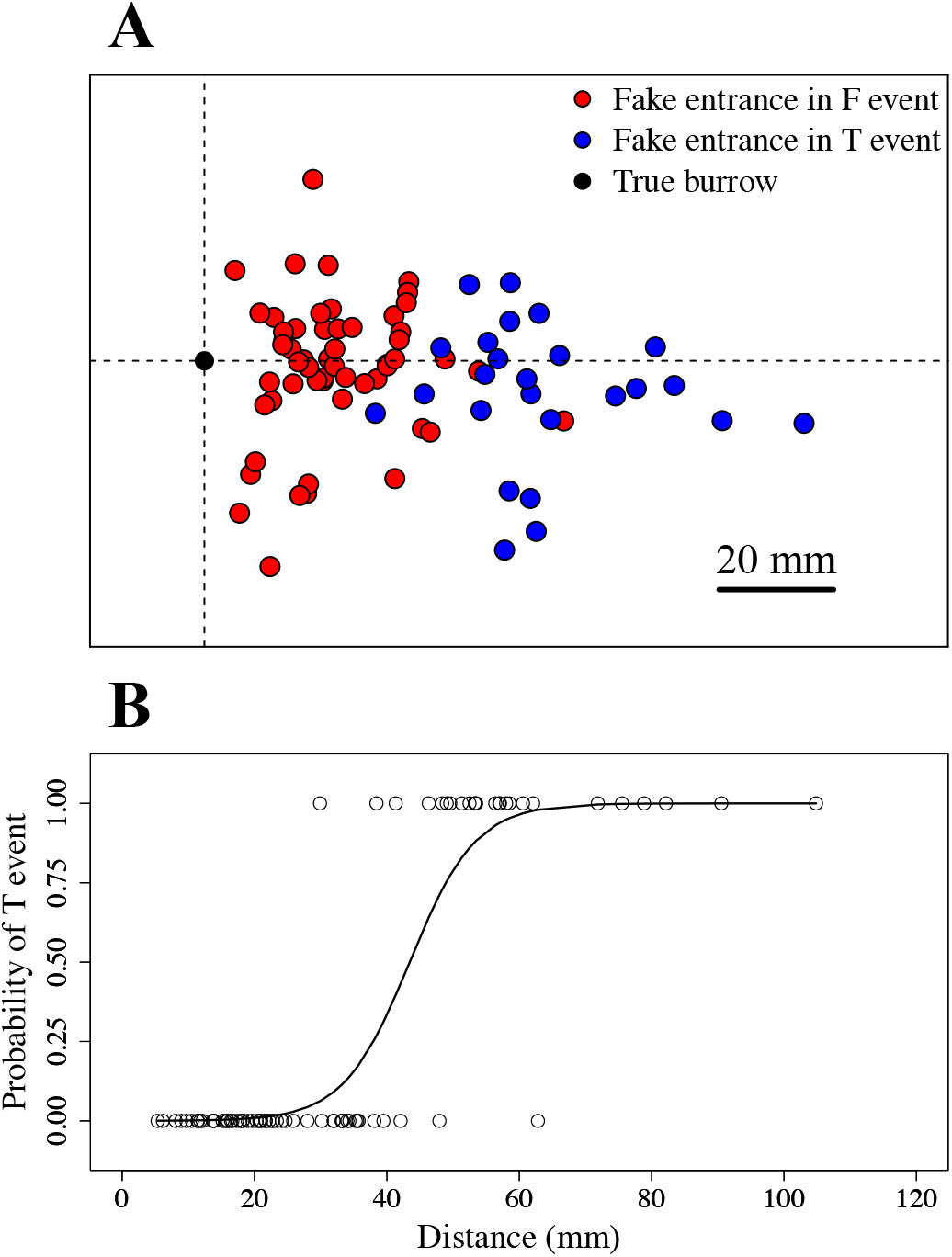
Burrow selection in relation to the distance between the true burrow and fake entrance under the non-social condition (experiments without territorial behavior). Data in this figure were adopted from Ref. [26] (CC BY 4.0) to allow direct comparison with the present results under the social condition (Fig. 2 in the main text). **(A)** Location distribution of fake entrances during F events (returned to fake entrance, red circles) and T events (returned to true burrow, blue circles). The black circle indicates the true burrow location. The direction from the starting point of the run to the true burrow is normalized to be right to left. **(B)** Probability of occurrence of a T event in relation to the distance between the true burrow and fake entrance. Open circles represent data from individual trials: T events (1) and F events (0). The solid line represents the best-fit sigmoid function.

